# The native structure of the Trichonympha centriole cartwheel reveals a zigzag stacking pattern

**DOI:** 10.64898/2026.04.08.717336

**Authors:** Carlee M. Rowsell, Shintaroh Kubo, Asuva Arin, Thibault Legal, Yining Yu, Khanh Huy Bui

## Abstract

Centrioles are essential cellular organelles with complex architectures. Their earliest assembly intermediate—the cartwheel—has a 9-fold symmetry established by Spindle assembly abnormal protein 6 (SAS-6). We used cryo-electron tomography and sub-tomogram averaging to resolve the native structure of the SAS-6 rings within the cartwheel of the exceptionally long *Trichonympha* proximal centriole. A 16-nm axial periodicity was formed by stacked rings of V-shaped SAS-6 tetramers, which formed the fundamental unit of the cartwheel’s central hub. SAS-6 head domains adopted a zigzag stacking pattern. Furthermore, the improved resolution provides new insights into the function of the previously observed central inner domain, which forms nine asymmetric densities that bridge adjacent tetramers, thereby imparting polarity and enhancing structural stability. Molecular dynamics simulations indicate increased rigidity in inter-tetramer interactions, supporting efficient ring formation and revealing how cartwheel architecture and polarity are established and stabilized.

## Introduction

Centrioles are microtubule-based, vital cellular organelles in eukaryotes (1), essential for establishing cell polarity, coordinating the cytoskeleton during mitosis, and forming cilia (2). Structurally, the centriole is roughly 250 nm in diameter and 500 nm long, and consists of nine triplet microtubules arranged with 9-fold radial symmetry (3). Upon docking in the plasma membrane via accessory structures and distal appendages (4, 5), a centriole becomes a “basal body” serving as the scaffold and template for cilia biogenesis (6, 7). Correct orientation of the centriole when docked onto the membrane relies on the centriole’s intrinsic polarity, such that its proximal end faces the cytoplasm, and its dynamic distal end engages connective fibers and appendages to anchor the organelle in its new role (*8, 9*). In keeping with its polar nature, the centriole’s internal architecture also varies along its length, with the proximal and distal regions containing distinct substructures. While the distal centriole seeds the formation of the transition zone by gradually adopting the architecture of the ciliary axoneme, the proximal region is defined by the presence of a ∼100 nm scaffold structure known as the “cartwheel” at the start of the assembly.

First identified nearly 70 years ago in the termite gut symbiont *Trichonympha* (*10*) the cartwheel consists of a ∼20–25 nm-diameter circular central hub (CH) that radiates nine spokes, each terminating in a pinhead that is anchored to a peripheral triplet microtubule (*3, 11, 12*). The cartwheel, with its distinct geometry, serves as a critical molecular template during early centriole assembly, functioning as a structural ruler and enforcing the 9-fold radial symmetry essential for proper centriole architecture (*13*). Though it is degraded upon centriole maturation in vertebrates and some other species, the cartwheel structure persists in protozoa, suggesting organism-specific requirements for maintaining the centriole’s characteristic architecture (*14*). The CH lies at the core of the cartwheel and is primarily built from Spindle assembly abnormal protein 6 (SAS-6), a highly conserved, ∼70-kDa protein (*11*). SAS-6 possesses a modular architecture with three key domains: an N-terminal globular head that mediates ring oligomerization, a central coiled-coil region that drives homodimerization, and a highly flexible, species-variable C-terminal domain (*15*). SAS-6 dimers associate via both their N-termini and coiled-coil domains to assemble stackable cartwheel rings, though the precise higher-order organization of these ring structures has remained unclear (*16*). Moreover, while SAS-6 was first characterized in *Caenorhabditis elegans* (*17, 18*) and later in *Leishmania* (*19*) *Drosophila* (*20*), and *Chlamydomonas* (*15, 21*), it has been deemed essential to establish the cartwheel architecture and enforce 9-fold symmetry (*21*), in addition to being crucial for centriole formation and cell division at large (*17, 22-24*). While some evidence suggests that 9-fold symmetric assembly can occur independently of the cartwheel, disrupting SAS-6 expression or function typically leads to irregular triplet microtubule numbers and structural defects (*25*), which can propagate from the proximal centriole to the cilium (*6, 21, 26-28*). Mutational work has identified key residues for proper SAS-6 oligomerization and higher-order cartwheel assemblies, which has been demonstrated as exploitable for altering the overall shape of the CH (*29*) and modulating the symmetry of the cartwheels (*30*). SAS-6 can self-assemble into stable dimers, tetramers, and rings *in vitro* (*19, 20, 31, 32*), supporting its role in coordinating the symmetric CH ring. While the mechanisms by which SAS-6 rings assemble, stack and achieve polarity during centriole biogenesis have been interrogated (*33*) and relevant residues for overall centriole viability/disease models have been identified (*6*), there lacks a full understanding of the proximal centriole in the context of the cell. As such, several fundamental questions about cartwheel organization in its native context remain unanswered: how are individual SAS-6 rings oriented and stacked along the proximal-distal axis to generate the periodic cartwheel structure? What molecular features determine cartwheel polarity and ensure unidirectional assembly? How does the characteristic axial periodicity observed in cartwheels arise from the arrangement of SAS-6 oligomers?

Prior cryo-electron tomography (cryo-ET) studies of the cartwheel of *Trichonympha* spp. have exploited the extended proximal centriole regions of these organisms to provide critical insights into cartwheel architecture such as the 16-nm repeating units and the presence of the *Trichonympha*-specific central inner domain (CID) inside the CH (3) (*11*). However, detailed characterization of the organization of the cartwheel is still unclear due to resolution constraints. Further investigation in the cartwheel of *Paramecium, Chlamydomonas, Naegleria* and humans (*11*) has shown diversity in cartwheel organization in particular the spoke region, though the thickness of the ring density in the CH allows these studies to suggest that the CH has a “double ring” organization of SAS-6 rings. Structural and mutational studies of *Chlamydomonas* SAS-6 coiled coil *in vitro* indicates that the asymmetric SAS-6 coiled-coil complex is important for stacking and leads to a proposed model of ∼10° rotation offset in the “doublet ring” to accommodate the asymmetric coiled-coiling seen in cartwheel spokes (*12, 34*). However, the simple “double ring” model cannot explain why the 8-nm double-rings prefer to stack in line with another 8-nm ring to form the CH instead of continuing the rotational-offset assembly to form a helical symmetric structure, which would be probably preferable. Moreover, given cryo-ET of *in vitro Chlamydomonas* SAS-6 rings also suggest a direct mode of single ring stacking (*12*), and the exact high-order organization of SAS-6 ring formation within the CH has yet to be fully elucidated, there appears to be a critical void in our understanding of centriole biogenesis. Here, we use high-resolution cryo-ET and sub-tomogram averaging (STA) to resolve the native architecture of the *Trichonympha* proximal centriole at a sub-nanometer resolution, revealing its stacking polarity and characteristic “zigzag” pattern of SAS-6 oligomers *in situ*. We also resolve the molecular organization of the 16-nm periodicity, identifying the CID as a key structural element that stabilizes the ring-formation interface.

## Results

### 16-nm cartwheel periodicity in *Trichonympha* arises from stacked 8-nm rings

To investigate centriolar structure, we used the Pacific dampwood termite, *Zootermopsis angusticollis*, whose hindgut is known to contain three main *Trichonympha spp*.: *T. campanula, T. collaris and T. sphaerica* (*35*). Centrioles were isolated from the *Trichonympha spp. (3*), vitrified, and imaged by cryo-ET (Fig. S1). The appearance of isolated centrioles from the three species was consistent across tomograms, suggesting significant structural homogeneity, with the *Trichonympha* centriole proximal region exhibiting well-defined, periodic substructures of triplet microtubules and cartwheels (Fig. 1A-C, Fig. S2, 3, Table S1), consistent with previous studies (*3, 36*). The cartwheel has a 16-nm repeat with the spokes extending towards the triplet microtubule (Fig. 1B, C).

**Fig. 1.**
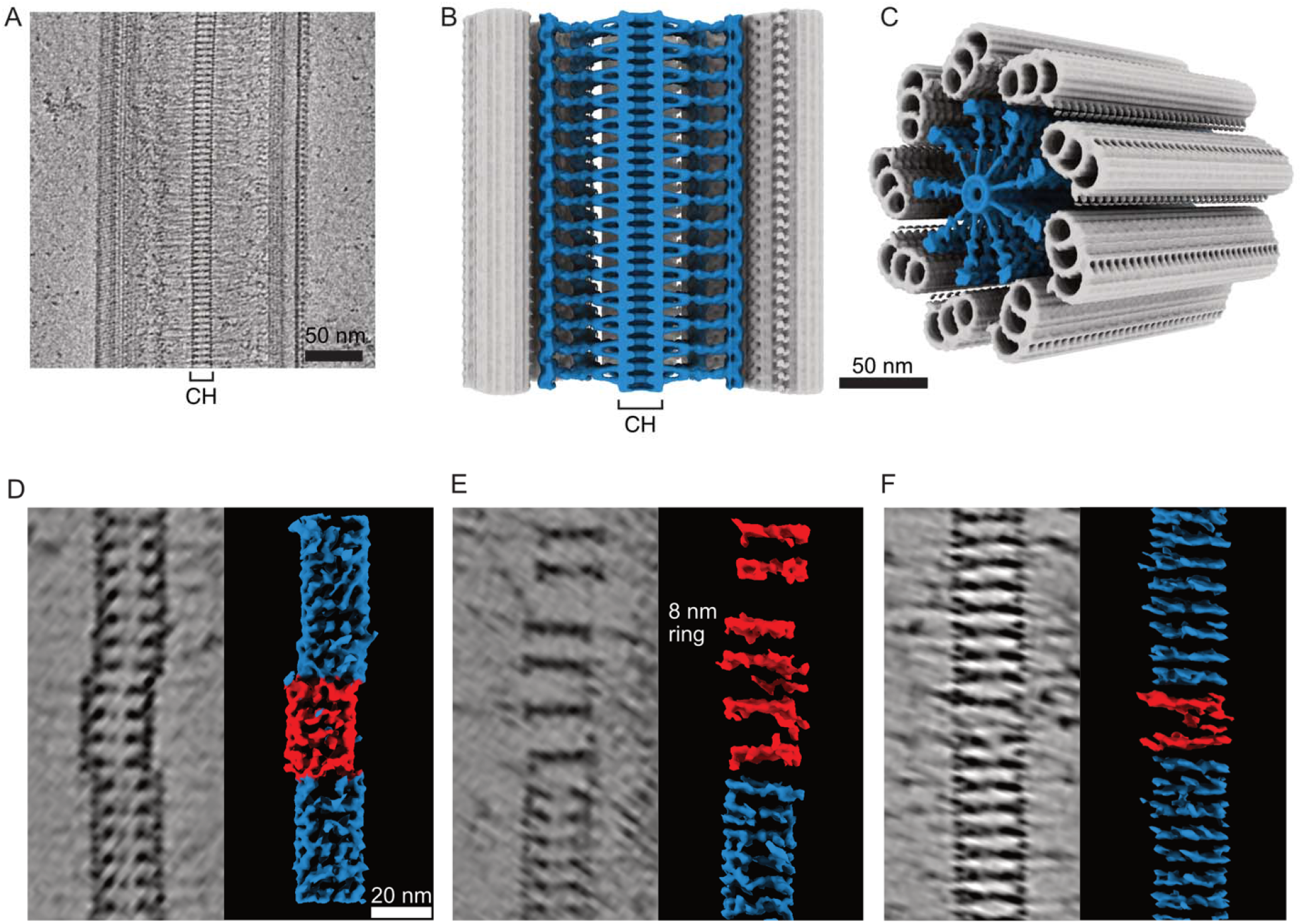
Architecture of the *Trichonympha* proximal centriole. **(A**) Tomogram snapshot of the CH. (**B**) Model of the longitudinal cross section of the proximal *Trichonympha* centriole constructed using averaged maps of the triplet microtubule (grey) and cartwheel (blue). (**C**) Alternate view of the proximal region. (**D–F**) Examples of non-intact cartwheels. The 8-nm ring appears as end-to-end Y-shaped densities. Tomographic slices and surface renderings of denoised and missing-wedge– compensated tomograms show: (**D**) 8-nm rings slipping out of register relative to adjacent rings; (**E**) separation of 8-nm rings, disrupting the canonical 16-nm axial periodicity; and (**F**) tilting of 8-nm rings relative to the CH axis. The regular region of the CH is shown in blue, whereas irregular 8-nm rings are shown in red.

The intact CH consists of 8-nm rings stacked along the central cartwheel axis. In tomographic slices, these rings appear as two Y-shaped densities joined end-to-end, forming a continuous column (Fig. 1A). Interestingly, our data contains cryo-tomograms of structurally perturbed cartwheels, likely from the purification steps. Partial disruption of the CH during sample preparation and/or vitrification shows individual 8-nm rings becoming laterally shifted or separated, with the most severe cases demonstrating full detachment from the stack of rings (Fig 1D-F). Detached rings frequently adopted oblique orientations relative to the overall axis, which directly exposed the modular organization of the overall assembly. These observations suggest that the 8-nm ring constitutes the fundamental cartwheel stacking unit, rather than the 16-nm repeat.

### SAS-6 forms a zigzag stacking pattern within the cartwheel hub

Comparing the STA maps of both 16-nm and 8-nm ring of the CH, the structural features in the 8-nm and 16-nm repeat maps of the cartwheel are only different at the spoke region (Fig. S3), confirming that the structure of the CH is faithfully captured at the 8-nm level. The 16-nm axial periodicity of the cartwheel arises from the stacking of two 8-nm rings, wherein spokes from successive 8-nm rings pair, align and associate to produce the 16-nm repeat which spans from the spoke to the triplet microtubules (Fig. 2A). Our measurements show that the CH repeating unit is 16.5 nm (Fig. S2B)-coinciding with the ∼16.45 nm periodicity observed for the triplet microtubules (Fig. S2B). For simplicity, we will refer to this CH periodicity as 16-nm throughout the paper.

**Fig. 2.**
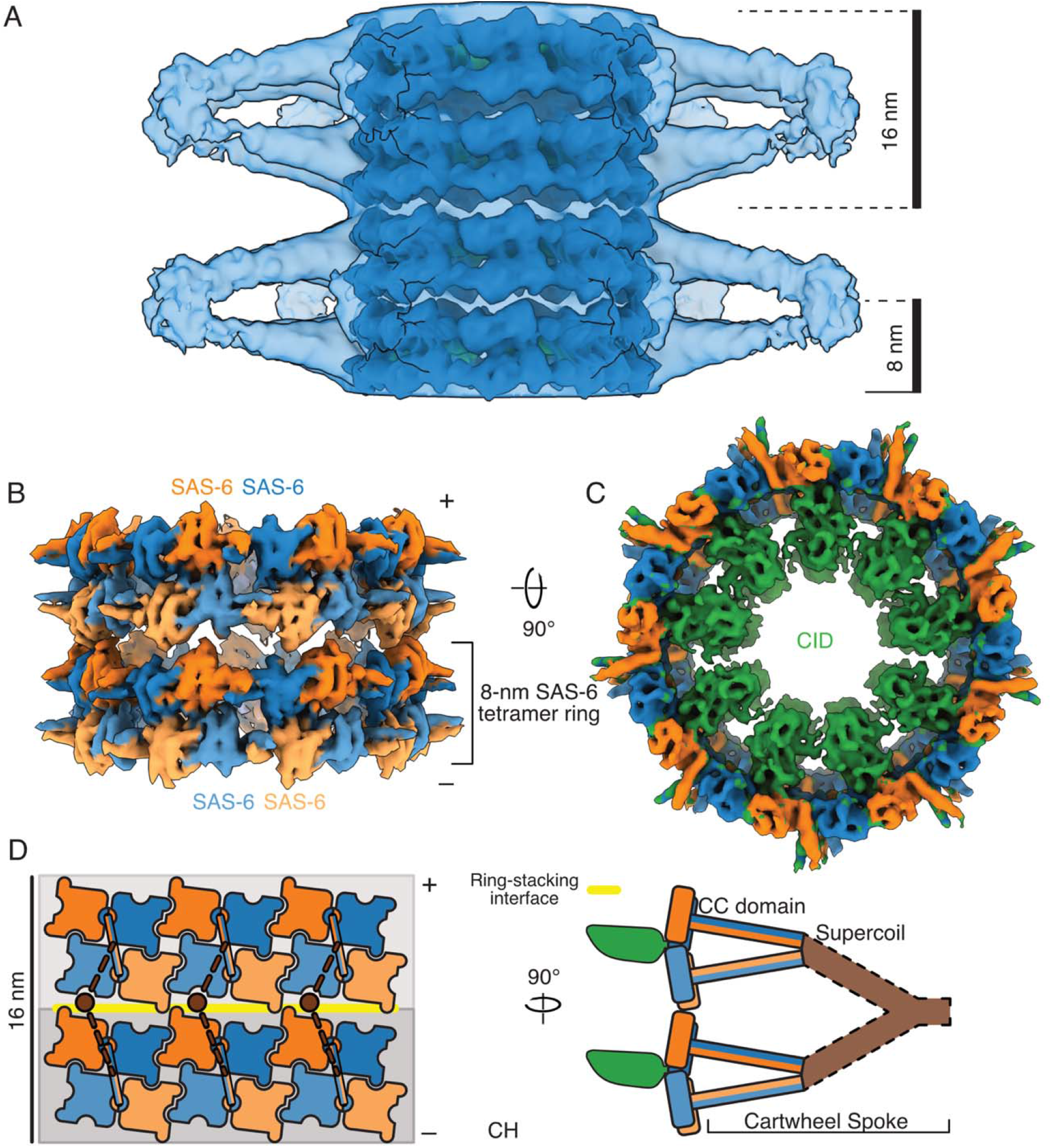
Molecular organization of the *Trichonympha* CH. (**A**) STA map of the 16-nm repeats of the CH. The dark blue surface is the map at 10.5 Å resolution, displayed at a high threshold to demonstrate CH organization. The transparent, light blue surface is the same map filtered at 20 Å and displayed at a low threshold to show the spokes. Two 8-nm ring that comprise each 16-nm repeating unit. (**B**) STA map of the 8-nm repeat of *Trichonympha* CH. Colored densities correspond to individual SAS-6 subunits. An 8-nm ring composed of tetramers constitutes the fundamental stacking unit. Symbols (+) and (–) signs denote the centriole’s distal and proximal ends, respectively. **(C)** Rotated view of the CH showing the CH ring plane and the CID densities (green) inside the CH. **(D)** Schematic of the 16-nm repeating unit of the CH formed by two 8-nm tetramer rings. Dimer subunits interact via the coiled coil (CC) domain. The CCs of two dimers form a supercoiled spoke, which extends to the triplet microtubule. CH tetramer rings stacking occurs vertically through a ring-stacking interface (yellow line), and their CCs dimerize to form the spoke (brown dotted line).

To resolve the architecture of the CH, we resolved the STA map of the symmetry-expanded 8-nm CH protomers to 7.6 Å resolution. Leveraging SAS-6’s high interspecies conservation (Fig. S4A), we used the existing *Trichonympha* SAS-6 sequence from *Trichonympha agilis* SAS-6 (TaSAS-6; UniProt ID: R4WPE9) to model the CH. Fitting the AlphaFold3-predicted structure (*37*) into the 8-nm map with DomainFit (*38*), which statistically assesses model placement, yielded a well-supported and confident molecular description of the CH (Fig. 2B, Fig. S4B-C, Movie S1). This also enabled the clear visualization of CID binding inside the CH (Fig. 2C).

Our fitting suggests that the fundamental building block of the 8-nm ring is a SAS-6 tetramer (Fig. 2D) and not the dimer, in spite of previous *in vitro* work demonstrating dimer-based SAS-6 ring formation (*15, 33, 34*). Each tetramer is composed of two SAS-6 dimers, which form a C2 symmetric subunit (Fig. 2D, purple color) (*20*). Together, the stacked tetrameric rings form the 16-nm repeat of the CH. This arrangement is consistent with the simple offset double ring model proposed previously from cryo-ET of centrioles of different species and the *in vitro* asymmetric complex of *Chlamydomonas* coiled coil regions (*11, 12, 34*).

### SAS-6 tetramer adopts a V-shape conformation

We wanted to further characterize the architecture of SAS-6 in the CH. In the zigzag pattern of the CH, each SAS-6 tetramer forms a parallelogram shape that continues laterally nine times to form an 8-nm ring (Fig. 3A, B). Regarding stacking, the SAS6 tetramer ring aligns in the same register as the neighboring ring, with an inter-ring distance of 82 Å along the cartwheel axis (Fig. S4E).

**Fig. 3.**
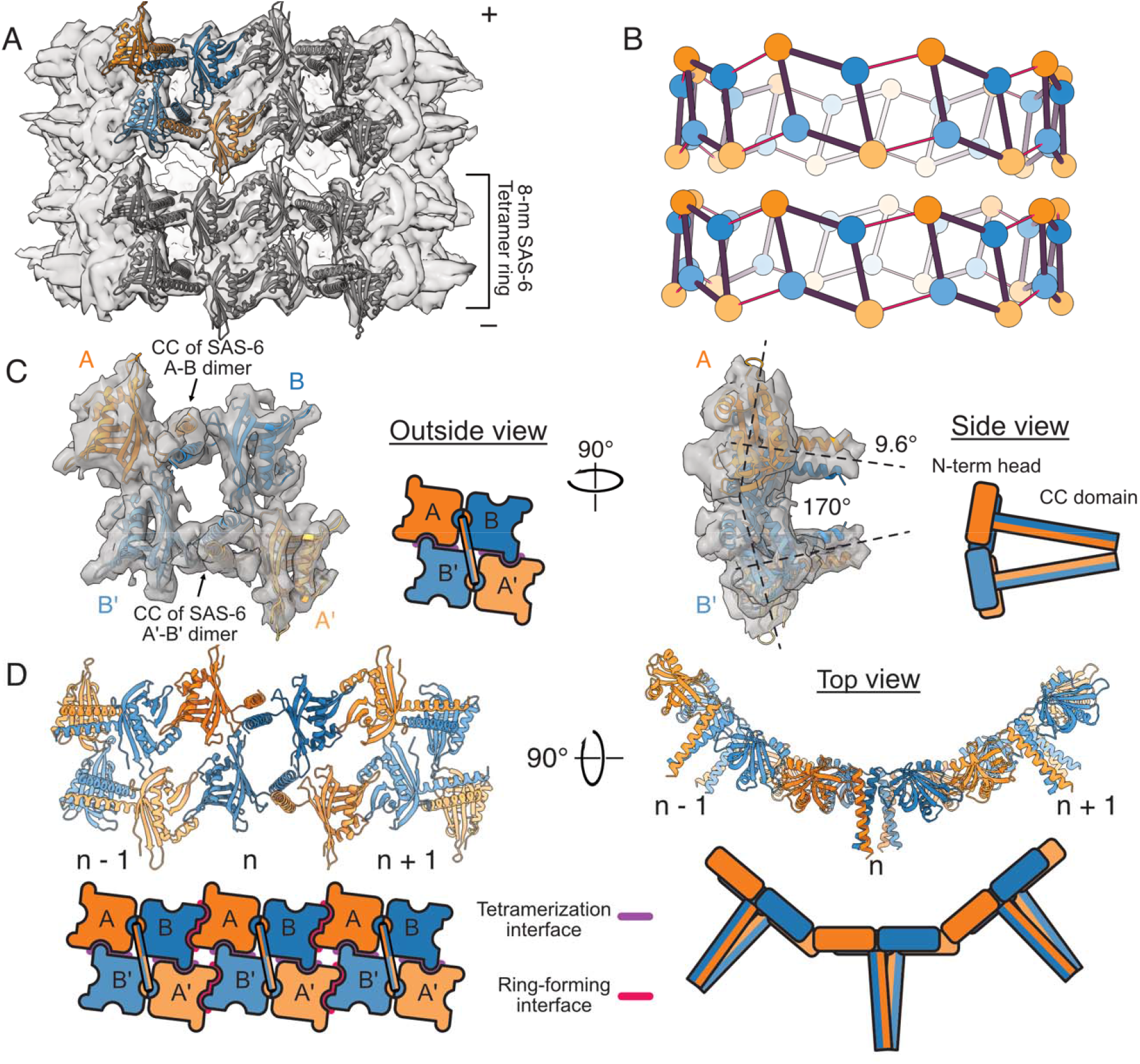
SAS-6 ring forms a zigzag stacking pattern within the cartwheel CH. (**A**) STA map of the 8-nm repeat of *Trichonympha* CH fitted with 4 SAS-6 tetramers. One tetramer is colored to distinguish four individual SAS-6 subunits while three others are colored in gray. The map shows two stacked 8-nm tetramer rings. Symbols (+) and (–) signs denote the centriole’s distal and proximal ends, respectively. (**B**) Zigzag stacking ring model of SAS-6 organization based on the fitting in (A). The center of mass of individual SAS-6 N-terminal head domains are displayed in colors consistent with (A) of asymmetric SAS-6 subunits within a tetramer (black parallelogram). Thin red lines indicate the tetramer-tetramer interaction within the same 8-nm tetramer ring. **(C)** Front (left) and side (right) views of a single SAS-6 tetramer fitted into the STA map. SAS-6 subunits A and B form one dimer. A′ and B′ form a second SAS-6 dimer, which pairs with the A/B dimer to form a tetramer in a C2-symmetric arrangement. The tetramerization interfaces between the A/B and A′/B′ dimers are indicated in purple. **(D)** Ring-forming interfaces (red) between neighboring SAS-6 tetramers within the 8-nm tetramer ring. Three neighboring tetramers (n-1, n, n+1) are shown.

Within a tetramer, the offset rotation of between the top and bottom SAS-6 dimers within the 8-nm ring along the CH center axis is calculated to be ∼6.7° (Fig. S4E). This rotational offset is underpinned by the tetramer’s unique geometry: the two SAS-6 dimers adopt a V-shape conformation of ∼170° along the dimer-dimer interface, facilitating the convergence and oligomerization of their coiled-coil regions into a supercoil within the same tetramer (Fig. 3C). The tetramerization interface involves helix α1 and loop β2–β3 of chain A′, and loops β1–β2 and β3–β4 of chain B and the same helix and loops of chain A and B’ (Fig. S5A-C). Notably, the V-shape tetramer conformation makes a distinct difference between the tetramerization interface (Fig. 3D, purple) and the ring-stacking interface (Fig. S5G). The tetramerization interface requires inward pointing SAS-6 dimers while the ring-stacking interface requires outward pointing SAS-6 dimers. That makes the tetramerization interface not available at stacking region of the 8-nm tetramer ring, allowing the 8-nm tetramer ring to stack at the same register onto each other. Earlier STA maps of other species show consistent features with our map of *Trichonympha* such as offset coiled coil in *Paramecium (11*) or tilted coiled coil in *Teranympha mirabilis* (*12*). This suggests other species have similar CH architecture as *Trichonympha*.

We also compared our model with the single-ring SAS-6 model from *Leishmania* SAS-6 crystal structure (*19*) (Fig. S5D, E). The dimer of *Leishmania* SAS-6 forms an angle of 3.6° with the ring plane while it is twice of that (7.3°) in our tetramer model. Furthermore, when superimposing the *Leishmania* SAS-6 crystal structure with our tetramer ring model, the SAS-6 N-terminal head of the adjacent dimer shows a large rotation of ∼23° along the ring plane (Fig. S5E) though still maintaining the lateral head-domain contacts seen in crystal structures of *Leishmania* and *Chlamydomonas* (*19, 32*) (Fig. S5F). This interface is likely to be mediated by Q73 residue (from one subunit) interacting with Y118 residue in TaSAS-6 (which corresponds to the conserved F118 position in other species) and R119 residue from the partner subunit (Fig. 3D, Fig. S5F). These contacts which are largely conserved (*19, 30*) act as a hinge for the interactions of two tetramers. Thus, though the SAS-6 tetramer retains the same ring-forming interface with previously known dimer structure, SAS-6 tetramer represents a unique conformation, which allows the zigzag model of V-shape SAS-6 tetramer.

The observation that isolated 8-nm rings persist in perturbed cartwheels suggests that the tetramer constitutes the fundamental unit forming individual 8-nm rings, which in turn serve as the basic stacking units of the CH. Consistent with this idea, *Drosophila melanogaster* SAS-6 has been shown to tetramerize both natively and when recombinantly expressed (*20*), supporting a tetramer-based model of cartwheel assembly. Crystal structure of *Chlamydomonas* SAS-6 reveals asymmetric interaction between two coiled coils (PDB: 6YRL) and mutation perturbed this asymmetric interface seems to interfere with *in vitro* ring formation (*34*). These results support that SAS-6 itself can tetramerize independently prior to the formation of the 8-nm ring (Fig. S5H).

### The CID confers cartwheel polarity and stabilizes the tetramer-ring interactions

Although the centriole is a polar organelle, our proposed model of the CH does not exhibit any intrinsic polarity. Prior work in *Trichonympha* reported that the CID is offset relative to the spoke-density axis, suggesting that it could impart or help maintain centriole polarity [17]. We unambiguously resolve the CID as nine asymmetric densities located within the lumen of the 8-nm tetrameric CH (Fig. 4A–B, Fig. S3, Fig. S6A). Each CID adopts an egg-shaped morphology with a prominent finger-like extension (Fig. S6B), providing a clear polarity marker.

**Fig. 4:**
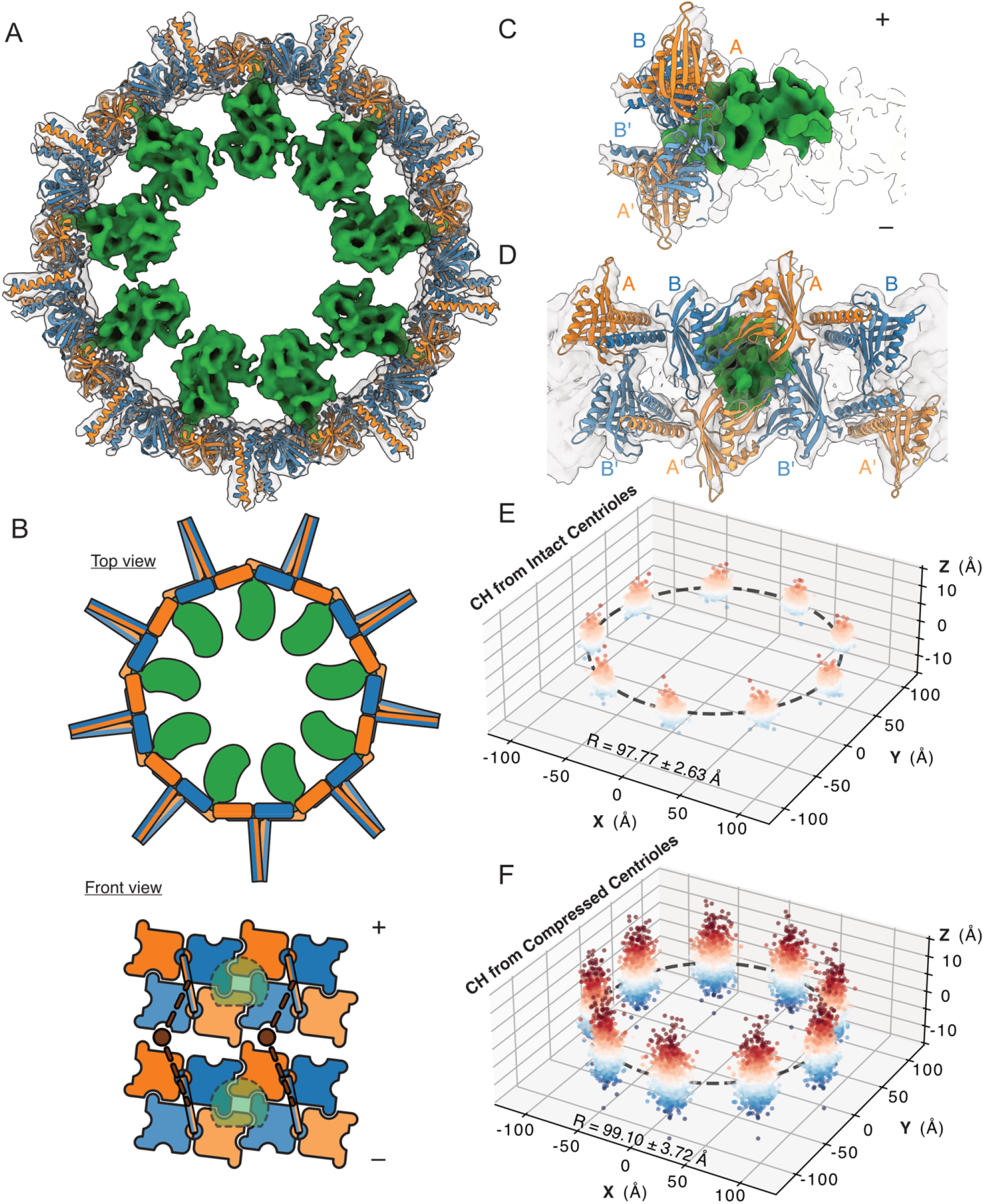
The CID density reinforces the inter-tetramer ring interface. **(A)** Cross section of the CH ring looking from the distal to proximal direction (CID, green; SAS-6, blue and orange; rendering threshold, 0.00267). **(B)** Cartoon models of the 16-nm CH unit with CIDs. **(C)** Orientation of the CID in the tetramer ring. Symbols (+) and (–) signs denote the centriole’s distal and proximal ends, respectively. **(D)** Orientation of the CID between SAS-6 tetramers showing the clear finger-like density inserting into the tetramer-tetramer interface. (**E, F**) 9-fold symmetrized three-dimensional plot of the coordinate of X, Y and Z-offset of the protomers of 8-nm rings of five intact centrioles (**E**) and five compressed centrioles **(F)**. The points are colored based on Z-offset values. The dashed circle denotes the estimated mean radius (R), which increased from 97.77 ± 2.63 Å in intact centrioles (n = 613 from 5 tomograms) to 99.10 ± 3.72 Å in compressed centrioles (n = 455 rings from 5 tomograms).

Remarkably, the CID bridges adjacent tetramers by binding at the tetramer–tetramer interface (Fig. 4C, D). Based on analysis of ring-formation interactions, the hydrophobic interaction between N-terminal domains is relatively weak (K_d_ for C. elegans, zebrafish, and human SAS-6s range from approx. 50– 110 μM) (*33*). The CID appears to significantly stabilize this interaction through its finger-like extension, which inserts into the junction of four SAS-6 molecules from two adjacent tetramers (Fig. 3D). At the molecular level, the CID contacts helix α1 (residues 63–73) of SAS-6 subunits B and B′, as well as loops β1–β2 (residues 15–21) and β5–β6 (residues 104–109) of subunits A and A′ (Fig. S6C– D). Because these interaction regions—especially loops β1–β2 and β5–β6 (Fig. S4A)—are not strongly conserved across species, CID function may be specific to *Trichonympha* spp.

Our map also suggests the presence of linkers connecting consecutive CIDs, forming a secondary inner ring approximately 14.7 nm in diameter (Fig. S6A). This inner ring may act as a structural scaffold that stabilizes the CH, explaining why the *Trichonympha* cartwheel is unusually long and regular compared to other species. Alternatively, the CID could serve as a nucleation point for assembling the SAS-6 tetramer ring, thereby promoting the formation of an extended cartwheel. Three-dimensional classification of the CID density on symmetry-expanded 8-nm subunit protomers (Fig. S6E) shows that the only 3.4 % of particles lacks CID, indicating high occupancy driven by strong binding or inter-CID interactions.

To test the robustness of this assembly, we analyzed the effect of centriole compression on the shape of the CH (Fig. S6F). In significantly compressed centrioles, protomer positions vary more within the ring plane compared to intact centrioles (Fig. 3E–F). However, the ring diameter increases only slightly, from 97.77 to 99.10 Å while the Z-offset of protomers from the ring plane increases from 0.69 to 1.67 Å, suggesting decreasing in ring planarity (Fig. S6G). Despite significant centriole compression, it is remarkable that the 8-nm ring remains round, provided the weak ring-formation interface from SAS-6 alone. These analyses consolidate our hypothesis that the CID significantly stabilizes the 8-nm ring against mechanical deformation in *Trichonympha*.

### SAS-6 tetramers facilitate ring formations

To investigate how SAS-6 rings assemble, we performed coarse-grained molecular dynamics simulations to understand whether ring formation is more efficiently nucleated by inter-tetramer interactions rather than by interdimer interactions. We set up simulations involving (i) two dimer subunits or (ii) two tetramer subunits corresponding to segments of the SAS-6 ring (Fig. 4A, Fig. S6).

These simulations revealed that the ring formation interface (between the dimer/tetramer interface) acts as the hinge for interaction (Fig. S7), as suggested by the difference between our model and crystal structures of SAS-6 from other species (Fig. S5C). In addition, tetramers are substantially more rigid than dimers in both their in-plane angle (within the ring plane) and their out-of-plane tilt (Fig. 4B–D, Fig. S7, Movie S2 and S3). This marked reduction in flexibility indicates that when SAS-6 subunits assemble through inter-tetramer interactions, they are far more likely to adopt and maintain conformations conducive to ring closure. In particular, the limited tilt variation in tetramers (Fig. 4C– D) favors a planar, closed ring rather than a helical or offset intermediate, supporting more efficient ring formation. SAS-6 assembles more efficiently on a surface, reminiscent of how the SAS-6 of the daughter centriole emerges orthogonally to a surface surrounding its mother centriole (*33*). This is consistent with our simulation, in which the out-of-plane tilt angle of the ring-forming interface is flexible and would be significantly constrained if SAS-6 assembled on a flat surface.

During the simulations, both dimers and tetramers fluctuate around angles associated with rings smaller than a perfect 9-fold ring, suggesting that SAS-6 assembly has an intrinsic propensity to form sub-9-fold intermediates. The structural position of the CID strongly suggests that it could lock the inter-tetramer angle, stabilize this ring-forming interaction, and enforce the bent geometry required for a canonical 9-fold ring and expected coiled-coil spoke assembly (Fig. 4E).

Together, these results support a model in which the cartwheel ring assembles directly from SAS-6 tetramers, which then stack through SAS-6 supercoil interactions to generate the 16-nm periodicity before further elaboration into the complete cartwheel. Moreover, in *Trichonympha*, the CID protein may stabilize the inter-tetramer interface, promoting the efficient nucleation of 9-fold symmetric rings, potentially explaining the organism’s unusually long proximal centriole regio. In other species, such as mice or humans, the densities observed within the cartwheel hub are less regular than the CID, suggesting that cartwheel-stabilizing factors may be species-specific.

## Discussion

Our reconstruction of the *Trichonympha* centriole proximal region with the clearly resolved CID and its asymmetric engagement at the inter-tetramer interface reveals a cartwheel architecture built of V-shaped SAS-6 tetramers forms the 8-nm ring units. Our data suggests that the 8-nm ring formation may be an intrinsic property of SAS-6 which allows these rings to form and stack on one another. Together, these observations suggest that the SAS-6 tetramer may represent a broadly used assembly motif, whose precise geometric implementation such as inter-coiled-coil angle, tilt, and stacking mode, is adapted in a species-specific manner, as previously observed for cartwheel coiled-coil packing (*19, 22*). Direct visualization of SAS-6 tetramers in *Drosophila* (*20*) supports the conservation of this assembly principle. In contrast, fluorescence correlation spectroscopy in human cells indicates that SAS-6 is predominantly dimeric in the cytoplasm (*39*), leaving open whether a tetrameric form exists prior to cartwheel incorporation or whether tetramerization is regulated in a species- or context-dependent manner.

The asymmetric positioning of the CID in *Trichonympha* provides a molecular polarity mechanism, preferentially engaging the distal SAS-6 dimer and coinciding with the altered conformations between distal and proximal dimers in the tetramer. Interestingly, densities inside the CH but not necessarily the same as CID in Tetrahymena are observed in CH of many species (*11, 12*). In organisms lacking a clearly defined CID, polarity may be imposed by intrinsic asymmetric features of SAS-6’s coiled coil interactions or distinct luminal factors (*e*.*g*., cartwheel regulators like STIL centriolar assembly protein) that bias one side of the hub (*3, 22*).

Our structural and simulation data support two non-exclusive roles for the CID. First, CID clearly stabilizes the inter-tetramer junction: its finger-like insertion and contact with the SAS-6 helix α1 and nearby loops could lock the inter-tetramer angles and reduce local flexibility, consistent with our molecular dynamic results indicating that constrained tetramers favor ring closure. Second, the CID could act as a nucleator by providing an inner scaffold that templates favorable ring geometry and promotes sequential ring stacking, an idea reminiscent of the inner-core scaffolds proposed in computational and experimental studies (*3, 40*). Monobody studies demonstrate that modest perturbations of SAS-6 architecture can shift assembly outcomes from rings to helices or other oligomers (*29*) indicating that luminal binders can both stabilize and re-shape SAS-6 assemblies in addition to some small molecule tuning data (*41*). Thus, CID could act both to lock geometry (as a stabilizer) and to bias nucleation toward canonical, vertically stacking 9-fold rings (as a polymerase) (*33*).

Our simulations suggest an intrinsic tendency for smaller, sub-9-fold intermediates without the CID, indicating that SAS-6 alone produces a range of curvatures. This is consistent with cell free *in vitro* assembly of *Chlamydomonas* SAS-6 N-terminal region (*29, 30*). Analysis of our ring diameters showed that ∼80% of ring assemblies were 9-fold symmetric, with ∼20% adopting eightfold symmetry (*29*). Therefore, CID-mediated locking of inter-tetramer angles could be a key enforcing mechanism that biases the assembly toward the canonical 9-fold geometry, consistent with crystallographic and *in vitro* assembly studies showing that accessory factors are needed to achieve robust, reproducible 9-fold cartwheels (*22, 32*). Our data suggests that the *Trichonympha* CID augments SAS-6’s intrinsic oligomerization propensity by restraining conformational freedom and promoting the biologically required cartwheel symmetry and length of the cartwheel.

Together, our findings define general principles by which SAS-6 tetramers and luminal factors cooperate to establish symmetry, polarity, and stable stacking during cartwheel assembly, allowing us to propose an assembly model (Fig. 5). In this model, SAS-6 tetramers assemble into 8-nm rings, a process promoted or stabilized by associated factors such as the CID in *Trichonympha*. Alternatively, there is a possibility that SAS-6 dimers assemble into 8-nm tetramer rings directly through associated factors. These tetrameric rings then stack longitudinally. The pairing of the resulting supercoils gives rise to the spokes that define the 16-nm repeating unit. This process of forming 16-nm repeating spoke might involve other proteins such as SAS-5, which forms large oligomers (*42, 43*) and is known to interact with SAS-6 (*23*) in addition to cartwheel assembly contributors Plk4/ZYG-1 and STIL (*44, 45*). Extending this framework to other species, including humans, will be essential to determine whether this mechanism represents a universal principle of centriole biogenesis.

**Fig. 5.**
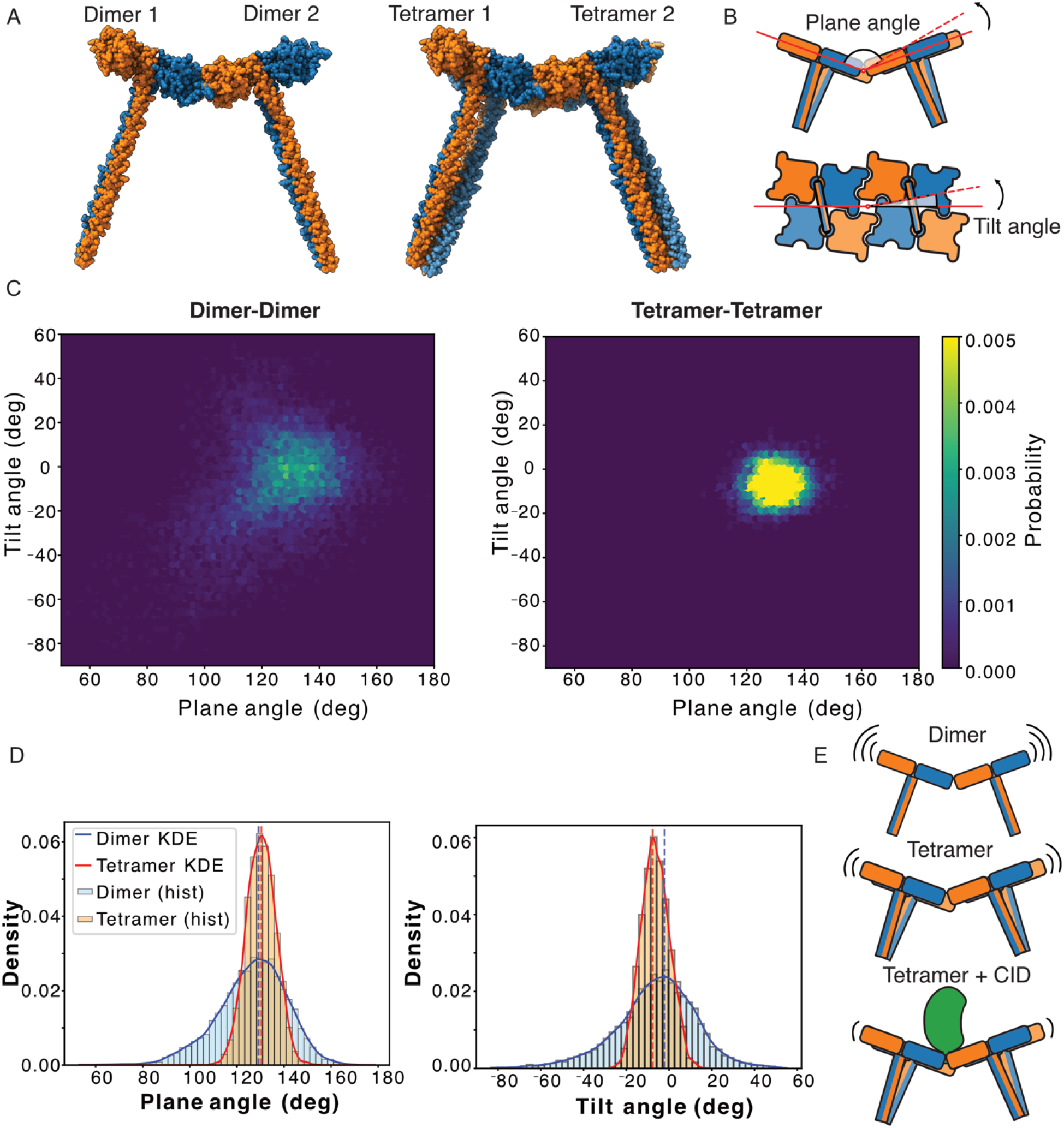
Coarse-grained molecular dynamics suggest that inter-tetramer interactions are more than those between dimers. (**A**) Starting models used for the interdimer and inter-tetramer simulations. (**B**) Schematic definitions of the plane and tilt angles used in our analysis. (**C**) Hexbin plots showing the joint probability density of tilt and plane angles for the interdimer *versus* inter-tetramer simulations. (**D**) Histograms with overlaid kernel density estimation (KDE) curves for the plane and tilt angle distributions in the two simulation conditions. For plane angle, 140° corresponds to the 9-fold symmetry ring and the starting conformation. (**E**) Proposed model of CH assembly. Interdimer interactions are highly flexible, whereas inter-tetramer interactions are more stable. When the CID binds at the inter-tetramer interface, the resulting complex stabilizes further, favoring the formation of a 9-fold symmetric ring.

**Fig. 6:**
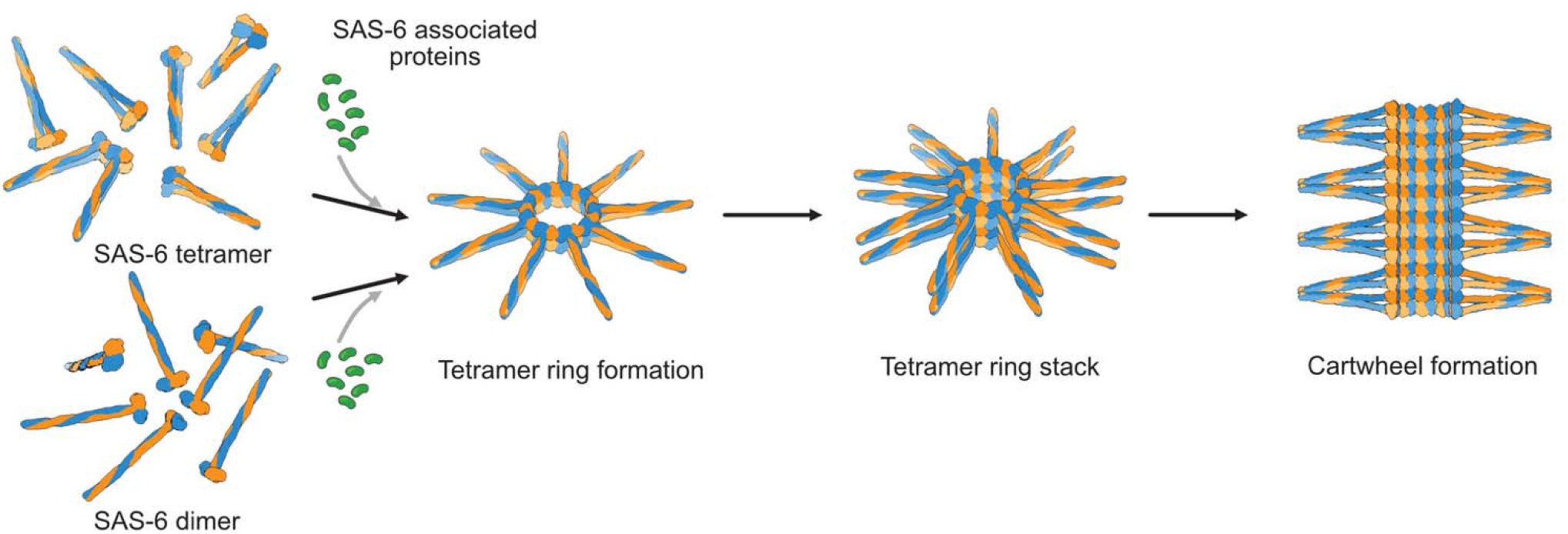
A proposed model of cartwheel assembly based on *Trichonympha* cartwheel structure.

## Acknowledgements

We thank Dr. Patrick Keeling (University of British Columbia) for generously providing the termite sample. We thank Drs. John Bergeron and Albert Berghuis and High-Fidelity Science Communications for manuscript editing. We thank Drs. Kaustuv Basu and Kelly Sears at McGill’s Facility for Electron Microscopy Research for help with data collection. K.H.B. is supported by grants from the Canadian Institutes of Health Research (PJT-190195) and the Natural Sciences and Engineering Research Council of Canada (RGPIN-2022-04774). T.L. is supported by the Fonds de Recherche du Québec – Santé (FRQS) fellowship (353841). A.A. is supported by the CRBS studentship and McGill internal studentship.

## Author contributions

Conceptualization: K.H.B. Methodology: C.M.R., A.A., S.K., and K.H.B. Investigation: C.M.R., A.A., S.K., T.L., Y.Y. and K.H.B. Visualization: C.M.R., A.A., S.K., and K.H.B. Funding acquisition: K.H.B. Supervision: K.H.B. Writing – original draft: C.M.R., S.K., and K.H.B. Writing – review and editing: C.M.R., A.A., S.K., T.L., and K.H.B.

## Data availability

EM maps have been deposited in the Electron Microscopy Data Bank (EMDB) under accession numbers: EMD-74424, EMD-74426, EMD-74421, EMD-74427, EMD-76289 corresponding to 8-nm and 16-nm CH maps, 8-nm and 16-nm CH subunit maps, and the triplet map.

The model for the fitted SAS-6 subunits in the 8-nm subunit map is deposited in the Protein Data Bank (PDB) under accession numbers: 9ZMK.

## Competing interests

The authors declare no competing interests.

## Notes

### Competing Interest Statement

The authors have declared no competing interest.

