## Supplementary Material for "The native structure of the Trichonympha centriole cartwheel reveals a zigzag stacking pattern"

### Materials & Methods

#### Trichonympha spp, isolation

*Zootermopsis angusticollis* termites were anesthetized on ice prior to hindgut extraction and placement in 10mM K-Pipes buffer (pH 7.2). *Trichonympha* cells (*T. campanula, T. collaris, and T. sphaerica*) were released from the gut upon rupture and left to sediment on ice for 10 minutes. Supernatant was aspirated and discarded, with the pellet resuspended in K-Pipes buffer. This sample washing was repeated three times to remove bacteria and other gut microbes.

#### Centriole purification

Isolated *Trichonympha spp.* cells were lysed following 1h standing incubation in 1 mL 10 mM K-Pipes 0.5% NP-40 and 1mM PMSF on ice, as per the protocol outlined in Guichard et al. 2013 (*1*). Following lysis, cell contents were pelleted via centrifugation at 500 x *g* for 3min at 4°C and supernatant was collected. Released centrioles and cilia were then pelleted via centrifugation at 1,000 x *g* for 5 min at 4°C and resuspended in TE buffer for EM sample preparation.

#### Negative staining and TEM

Isolated centrioles were placed on negatively glow-discharged (30mA, 1 minute) 300 mesh carbon film grids (Electron Microscopy Services #CF300-Cu-50), incubated for 1 minute, and blotted with filter paper (Whatman no. 1). Grids were then washed and blotted with water prior to a final single wash and 1 minute incubation with 2% depleted uranyl acetate (Electron Microscopy Sciences), after which they were blotted and let dry. Samples were then transferred to a Tecnai G2 Spirit Twin 120 kV Cryo-TEM (FEI) equipped with an AMT NanoSprint15 MK2 CMOS Camera and imaged at various magnifications with proprietary AMT software.

#### Cryo-ET sample preparation

Samples were mixed with 5nm gold beads with a 7:5 ratio and applied to negatively glow-discharged (30mA, 30s) Quantifoil 300 mesh R2/1 holey carbon copper grids (Electron Microscopy Sciences) inside the Vitrobot MkIV (Thermo Fisher) chamber. 5 microlitres of sample were incubated on the grid for 45 seconds at 100% and room temperature, then blotted with force 0 for 8 seconds and plunged frozen in liquid ethane.

Without cross-linking, the sample is flattened though the CH is intact. We used this dataset for STA of the 8-nm repeat. However, due to flattening of the basal body, the spokes did not keep good orientation in this sample. We prepared another sample cross-linked by glutaraldehyde (final concentration of 0.15%) for 30 minutes on ice and quenched by 1M Tris. This sample clearly improves the intactness of the basal bodies allowing us to obtain the 16-nm repeat of the cartwheel. STA of both datasets revealed no difference in CH structure.

#### Cryo-ET acquisition and reconstruction.

Dataset 1 (non-crosslinked, 62 tilt series) and dataset 2 (crosslinked, 121 tilt series) were collected via a Titan Krios 300 kV FEG electron microscope (Thermo Fisher) equipped with a direct electron detector K3 Summit (Gatan, Inc.) and a BioQuantum energy filter (Gatan, Inc.) using SerialEM (*2*). Acquisition was performed using a dose-symmetric scheme from -60 to 60 degrees in 3-degree increments at defocus values ranging from -1.5 to -3 μm. Micrographs were collected at a pixel size of 2.12 Å with a total dose of ~120 e− per Å^2. Preprocessing and CTF estimation were performed using the Warp pipeline (*3*), and tilt series alignment was completed using IMOD (*4*).

### Subtomogram Averaging

#### SAS6 STA

The CH subtomograms were manually picked using IMOD's 3dmod program, tracing through the centre of the cartwheel from proximal end to distal end based on triplet microtubule polarity determination. Subtomogram averaging was performed using Dynamo (*5*) following the filament-processing workflow(*6*). Tomograms were first Fourier binned to a pixel size of 14 Å/pixel, and subtomograms were interpolated at a periodicity of 80.56 Å and extracted in a 64-voxel box. In preparation for processing with RELION 5(*7*), particle coordinates and parameters were converted to STAR format with Warp, following which subtomograms were and re-extracted at 4.24 Å/pixel and of size 164 voxels, respectively. For further refinement, particles were imported into RELION for initial 3D refinement with C9 symmetry. Following classification to remove junk particles, sub-tomograms were 9-fold symmetry expanded, re-extracted in Warp at 3.53 Å/pixel using a 96-voxel box, and 9-fold symmetry expanded for another two iterations of 3D and refinement with C1 symmetry and tighter masking to one protomer. Refinement of the 8 nm unit was performed with 6523 particles, symmetry expanded to 58707, with a final round of un-binned M refinement (*3*) at 2.12 Å/pixel yielding a final resolution of 7.6 Å.

Processing of the 16-nm repeats began with 5594 particles extracted at 3.53 Å/pixel with a 96-voxel box. Prior to rescaling, these subtomograms were shifted by 42.4 Å. Particles were then symmetry-expanded around the cartwheel spokes (20° offset from the tetramer interface used for 8nm refinements), generating 50346 particles for subunit refinement, where final M refinements yielded a resolution of 10.98 Å.

#### Triplet STA

After tomogram reconstruction, triplet microtubules were manually picked using IMOD by tracing through the centre of the central B-tubule. Subtomograms were extracted with a 72-voxel box every 16 nm along these microtubules using a set of Dynamo scripts for microtubule alignment. Each subtomogram was aligned and averaged, using an initial reference generated from (EMD-42776) (*8*).

Following conversion to Relion-compatible STAR files with Warp, the resulting subtomograms were exported to Relion 5 at a pixel size of 6.30 Å/pixel. To obtain an intact triplet microtubule structure and enhance local structural details, focused classification and refinement were performed on the A- and C-tubule sequentially, using two region-specific masks: one mask targeted the disrupted protofilaments of the A-tubule, while the second mask focused on the broken protofilaments of the C-tubule. 4443 and 1117 particles were selected for the A and C tubule, respectively, yielding a final overall resolution of 15.2 Å for the microtubule triplet microtubule.

### SAS-6 Sequence Alignment

AlphaFold prediction of TaSAS-6 dimers shows retention of the conserved structure from known species. Multiple sequence alignment of SAS-6 homologs (*Chlamydomonas* UniProt: A9CQL4, *T. vaginalis* UniProt: A2G2L7 Human UniProt: Q6UVJ0, *T. agilis* UniProt: R4WPE9) using JalView (*9*). Sequences were truncated to encompass the N-terminus to the end of the first alpha-helical portion of the coiled-coil domain and aligned with Clustal Omega (*10*).

#### Visualisation

Tomograms were denoised using DeepDeWedge (*11*) for visualisation. Map post-processing with EMReady was performed for subtomogram-averaged maps at resolutions higher than 10 Å to improve interpretability (*12*). ChimeraX was used for the visualisation of maps and models (*5*). Replacement of subtomogram averages into the tomograms for visualisation as seen in Fig 1, was performed using subtomo2chimera.

#### Fitting of SAS6

For SAS-6 fitting, we used DomainFit (*13*) to fit the AlphaFold (*14*) predicted dimer of SAS-$6_{1\text{-}160}$ into the 8-nm subunit map using global search and to produce fitting statistics (20000 placement positions at a resolution of 7.6 Å). The top 16 positions reflect various positions of the SAS-6 dimer in the map and the C2 nature of the dimer (Fig. S4C). Performing the z-transform of the ChimeraX correlation scores, we calculated the hit p-values. The 16 top hits all have a z-score significantly separated from the rest (Fig. S4D) and have a p-value of $\sim 10^{-14}$, confirming they are true hits.

#### Model Building

The AlphaFold-predicted TaSAS-6 dimer was used as the initial atomic model, which were density-fitted as detailed above. Two SAS-6 tetramers were then rigid-body fitted into the map upon within ChimeraX to generate a final tetramer-tetramer arrangement. To accurately model the coiled-coiling of SAS-6 alpha helices, and the meeting of adjacent dimers' coils to form the supercoiled spoke, these helical domains were manually cut and repositioned in ChimeraX and reattached to their respective N-terminal domains using Coot (*15*). This model was then real-space refined using Phenix (*16*). To emulate the expected rigidity at the central hub interface due to spoke supercoiling for molecular dynamics simulations, artificial crosslinks were introduced between alpha-helices of the coiled-coil domain by creating cysteine bridges using Coot.

#### Angle and distance analysis of SAS-6 tetramers

All geometric analyses were performed relative to the 8-nm tetramer ring plane, defined as the XY plane, with the ring central axis along Z.

**Dimer tilt angle.** The angle of each SAS-6 dimer relative to the ring plane was measured as the elevation angle between the XY plane and the line connecting the centers of mass of the two SAS-6 N-terminal head domains (residues 1–130 of subunits A and B).

**Upper/lower dimer offset.** The rotational offset between the upper and lower SAS-6 dimers within the tetramer was measured as the rotation angle around the ring central axis (Z axis) formed by the centers of mass of the N-terminal heads of subunits A and B'. The ring central axis was determined by applying the ChimeraX command `measure rotation` to two consecutive tetramers. The ring diameter was calculated as the mean distance from the ring central axis to the centers of mass of the SAS-6 N-terminal heads.

**Inter-subunit rotation.** Rotational relationships between subunits within the tetramer were quantified using the ChimeraX command `measure rotation`, which returns the rotation matrix between any two subunits. Each matrix was decomposed into Euler angles representing rotations around the X, Y, and Z axes independently.

**Coiled-coil tilt angle.** The angle of each coiled coil relative to the ring plane was measured as the elevation angle between the XY plane and the line connecting the center of mass of residues at the coiled-coil start (residues 134–138 of subunits A and B) to the center of mass of residues at the coiled-coil end (residues 164–168 of subunits A and B).

**Inter-tetramer distance.** The axial distance between adjacent 8-nm rings was measured as the distance between the centers of mass of subunit A in two neighbouring tetramers. This line runs parallel to the ring axis (Z axis).

**Intra-tetramer vertical rise.** The vertical offset between subunits A and B' within the same tetramer was calculated by projecting the line connecting their centers of mass onto the ring axis (Z axis).

### CH ring offset analysis

The CH ring Z-offset analysis (Fig. 4E, F and Fig. S6F) was performed using Python scripts. To analyze the structural parameters of the nine-fold symmetric rings, raw particle coordinates (x, y, z) were extracted from the RELION-formatted STAR file. To account for arbitrary ring orientations within the tomographic volume, a local coordinate system was established for each ring. The coordinates were first centered at the geometric mean of the nine protomers within a single ring. Subsequently, Principal Component Analysis was performed on the centered coordinates, where the third principal component, representing the direction of least variance, was defined as the ring normal. A rotation matrix was calculated and applied to align this ring normal with the global Z-axis. To ensure a consistent reference frame across all rings, a secondary rotation around the Z-axis was applied to align the first protomer with the X-axis.

To account for the inherent nine-fold rotational symmetry of the protomer rings and to improve the signal-to-noise ratio of the spatial analysis, a rotational expansion procedure was applied to the centered and aligned coordinates. For each validated ring containing nine protomers, the dataset was augmented by rotating the original coordinates around the Z-axis in increments of 40°. This approach ensures that the resulting density maps and Z-deviation profiles reflect the cumulative spatial distribution of all protomer positions within the symmetry group, rather than being biased by the initial orientation of a single ring. The expanded coordinate set was then utilized to Z-offset heatmaps, providing a comprehensive visual representation of the ring's average radial consistency and axial planarity across the experimental groups.

Following alignment, structural parameters were calculated for each protomer to quantify ring dimensions. The radial distance (R) for each protomer was calculated as the Euclidean distance from the ring center in the XY plane ($R = \sqrt{x^2 + y^2}$). The Z-offset, or axial planarity, was defined as the displacement of each protomer from the calculated ring plane along the Z-axis ($Z_{offset} = z_{aligned}$). To assess the overall axial spread or "thickness" of the ring, the absolute Z-offset ($|Z|$) was utilized for quantitative comparison.

Quantitative comparisons between experimental groups were performed using a dual-track framework. Global distributions of radii and Z-offsets were visualized using Kernel Density Estimation and violin plots to identify structural subpopulations or shifts in planarity. To determine if structural differences between groups were statistically significant, an independent two-sample t-test was applied to the pooled protomer measurements for both radius and absolute Z-offset. All results were reported as mean ± standard deviation, with $p < 0.05$ considered statistically significant.

### Molecular Dynamics Simulation

#### Coarse grain molecular dynamic simulations

The starting models for MD simulation were constructed from SAS-$6_{1\text{-}250}$ AlphaFold3 dimer prediction. The dimers were then built into the dimer-dimer complexes or two tetramer-tetramer complexes based on our map. AlphaFold3 prediction of supercoil of SAS-6 tetramer was not working. Therefore, to simulate the coiled coil interaction of dimer's coiled coil within the same tetramer subunit, we mutation residues 235, 238 and 239 into cysteine in chain B and B'. These mutations lead to three-disulfide bonds (B 235 to B' 235, B 238 to B' 239, B 239 to B' 238) (Fig. S7A).

With that we have dimer-dimer complexes and tetramer-tetramer complexes (Fig. S7A) (i) two dimers interaction or (ii) two tetramer interaction for the MD simulations. In all simulations, the left-side chain A in Fig. S7A—was anchored at Cα of three residues: N63, Q73, and Q127. For both the dimer and tetramer SAS-6 systems, 10 independent simulations were performed. All MD simulations were conducted using CafeMol version 2.1(*17*), and each run consisted of $10^8$ MD steps. Underdamped Langevin dynamics at 300 K were employed. The friction coefficient was set to 0.02 (CafeMol units), and default parameters were used for all other settings. For both intra- and inter-molecular interactions, the AICG2+ force field, electrostatic interactions, and excluded-volume effects were included.

#### Calculation of plane angle and out-of-plane tilt angle

To quantify the relative motion of three reference residues during the simulation, we computed two geometric angles, plane angle and tilt angle, for every frame of a multi-model PDB trajectory. All analyses were performed with an in-house Python script.

For each frame, the Cα coordinates of three user-specified residues, $(\boldsymbol{r_1}, \boldsymbol{r_2}, \boldsymbol{r_3})$ = (T125 of Chain-A, Q127 of Chain-B, Q127 of Chain-C), were obtained from the PDB. From these coordinates, three vectors were defined:

$$\boldsymbol{v_1} = r_1 \to r_2, \qquad \boldsymbol{v_2} = r_1 \to r_3, \qquad \boldsymbol{v_3} = r_2 \to r_3$$

##### Plane angle

The plane angle was defined as the geometric angle between $\boldsymbol{v_1}$ and $\boldsymbol{v_2}$:

$$\theta_{plane} = \cos^{-1}\left(\frac{\boldsymbol{v_1} \cdot \boldsymbol{v_2}}{|\boldsymbol{v_1}||\boldsymbol{v_2}|}\right)$$

This angle describes the instantaneous opening angle formed by the three residues.

##### Reference plane construction

To evaluate out-of-plane motion, a reference plane was constructed using the initial configuration. The plane normal vector was defined as:

$$\boldsymbol{n_0} = \boldsymbol{v_1}(\boldsymbol{0}) \times \boldsymbol{v_2}(\boldsymbol{0})$$

where $\boldsymbol{v_1}(\boldsymbol{0})$ and $\boldsymbol{v_2}(\boldsymbol{0})$ are vectors from frame 0.

##### Tilt angle

The tilt angle (out-of-plane angle) measures how much the current $\boldsymbol{v_3}$ deviates from this reference plane. For each frame, $\boldsymbol{v_3}$ was decomposed into:

- a component parallel to the reference plane
- a perpendicular component along $\boldsymbol{n_0}$

The tilt angle was then computed using:

$$\theta_{tilt} = tan^{-1}\left(\frac{\boldsymbol{v_3} \cdot \boldsymbol{n_0}}{|\boldsymbol{v_3} - (\boldsymbol{v_3} \cdot \boldsymbol{n_0})\boldsymbol{n_0}|}\right)$$

which yields a signed tilt angle within ±90° indicating the direction of deflection.

In addition to the above calculation, we also analyse different ways to measure tilt and plane angles following only $\boldsymbol{v_3}$ or $-\boldsymbol{v_1}$ to see if the changes in tilt and plane angles are the direct results of flexibility at the hinge of dimer/dimer or tetramer/tetramer. In this case, the plane angle is measured as deviation from the original vector $\boldsymbol{v_3(0)}$ or $-\boldsymbol{v_1(0)}$. The tilt angle is measured by how much the vector deviates from the reference plane. In this case, the normal vector for the reference plane is defined as the cross product of $\boldsymbol{v_3(0)}$ and vector connect $\boldsymbol{r_2}$ and anchor point Cα of residues N63.

### Supplementary Movies

**Movie S1.** A representative tomogram, map, and structure of the *Trichonympha* centriole's central hub.

**Movies S2 and S3.** Representative trajectories for the interdimer (Movie S2) and inter-tetramer simulations (Movie S3). For each simulation setup, a portion of a representative trajectory was converted into a movie. Movies were compiled from 300 snapshots. Each snapshot was rendered using VMD. To create smoother visuals, VMD rendered the average of 5-frame segments. The frames were compiled into videos using ffmpeg and a sampling rate of 1 frame = 50,000 MD steps.

**Table S1**: Cryo-EM data collection and refinement statistics for all datasets used in this study.

| | Cryo-ET & Subtomogram averaging | |
|---|---|---|
| **Dataset** | **Dataset 1 (*Trichonympha* Centriole )** | **Dataset 2 (*Trichonympha* Centriole )** |
| **Microscope** | Titan Krios | Titan Krios |
| **Electron Detector** | Gatan K3 | Gatan K3 |
| **Zero-loss filter (eV)** | 20 | 20 |
| **Magnification** | 42,000 | 42,000 |
| **Voltage (keV)** | 300 | 300 |
| **Electron exposure (e/Å^2)** | 120 | 120 |
| **Defocus range (μm)** | 1.5-3 | 1.5-4 |
| **Pixel size** | 2.12 | 2.12 |
| **Tilt range (increment)** | -60° - 60° (3°) | -60° - 60° (3°) |
| **Tilt scheme** | Dose-symmetric | Dose-symmetric |
| **Tilt series acquired** | 62 | 121 |
| **Repeat unit (nm)** | 8 | 16 |
| **Symmetry imposed** | | |
| Ring | C9 | C9 |
| Subunit | C1 | C1 |
| **Subtomograms averaged** | | |
| Ring | 6523 | 5594 |
| Subunit | 58707 | 50346 |
| **Map resolution (Å)** | | |
| Ring | 10.9 | 12.2 |
| Subunit | 7.6 | 8.7 |
| **Refinement Statistics** | | |
| **Model** | **SAS-6 Central Hub** | |
| **Model-to-Map fit, CCmask** | 0.70 | |
| **All-atom clashscore** | 21.08 | |
| **Ramachandran plot** | | |
| **Outliers (%)** | 0.00 | |
| **Allowed (%)** | 2.53 | |
| **Favored (%)** | 97.47 | |
| **Rotamer outliers (%)** | 0.00 | |
| **Cbeta deviations (%)** | N/A | |
| **Cis-proline/general (%)** | 0.00 | |
| **Twisted proline/general(%)** | 0.00 | |

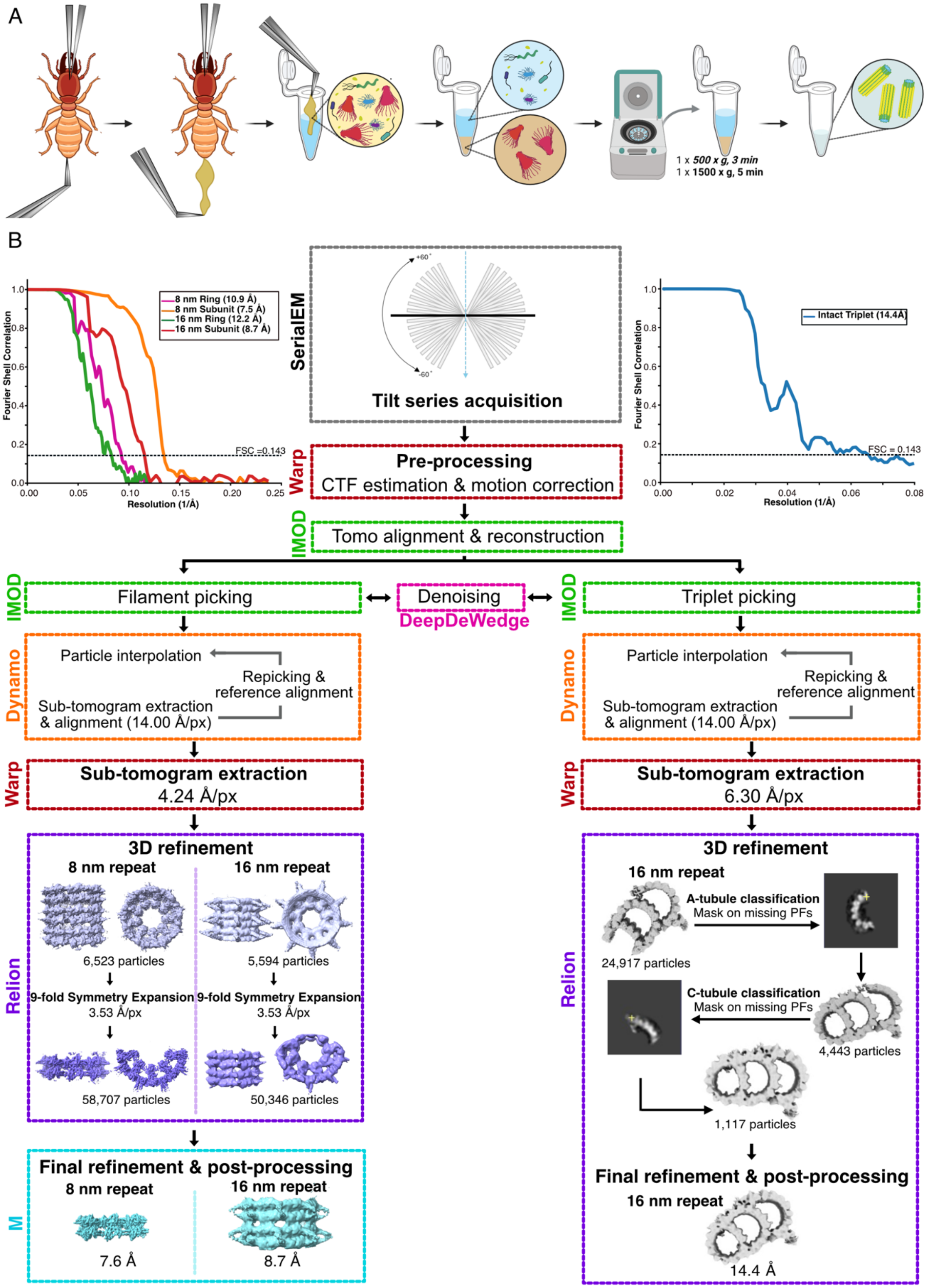


**Fig. S1: Centriole purification and cryo-ET & STA workflows.** (**A**) *Trichonympha* centriole isolation scheme from Pacific dampwood termite hindguts. Part of illustration from NIAID NIH BioArt Source. (**B**) Workflows used for cryo-ET STA of the cartwheel CH (left) and triplet microtubule (right), complete with gold-standard Fourier Shel Correlation (FSC) plots. The cartwheel FSC plot shows curves for the 8-nm and 16-nm rings and subunits. See Methods for details.

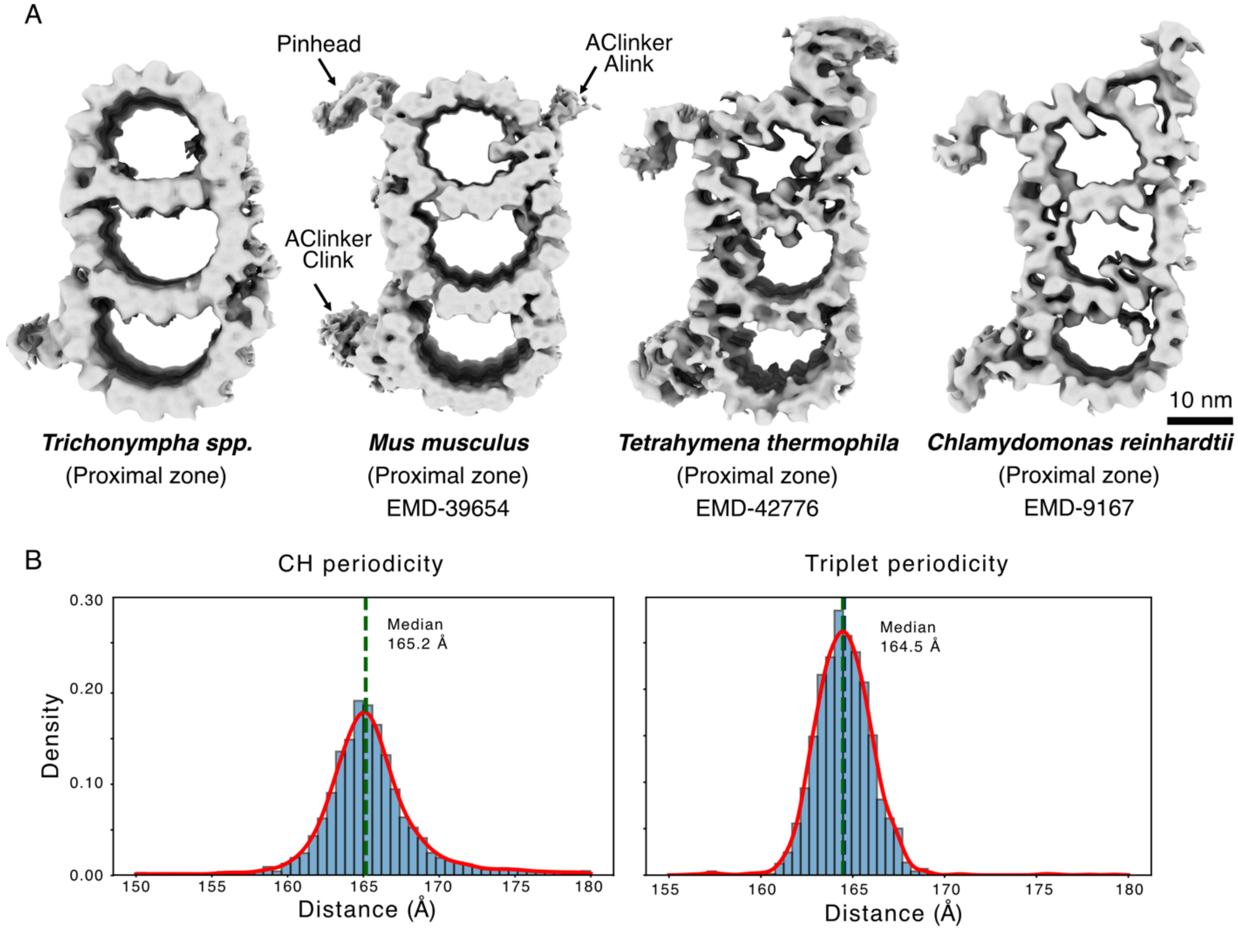


**Fig. S2**: **Comparison of triplet microtubules across species.** (**A**) Surface renderings of triplet microtubules from *Trichonympha* spp (this study), *Mus musculus* (EMD-39654) (*18*), *Tetrahymena thermophila* (EMD-42776) (*8*) and *Chlamydomonas reinhardtii* (EMD-9167) (*19*). All maps were low pass filtered to 15 Å for consistent visualization. The absence of the pinhead and AC-linker in the *Trichonympha* triplet microtubule reflects partial loss of these structures in the non-intact triplets captured in this dataset. (**B**) Periodicity analysis of the CH and the triplet microtubules, measured as the distances between consecutive particles in the tomograms.

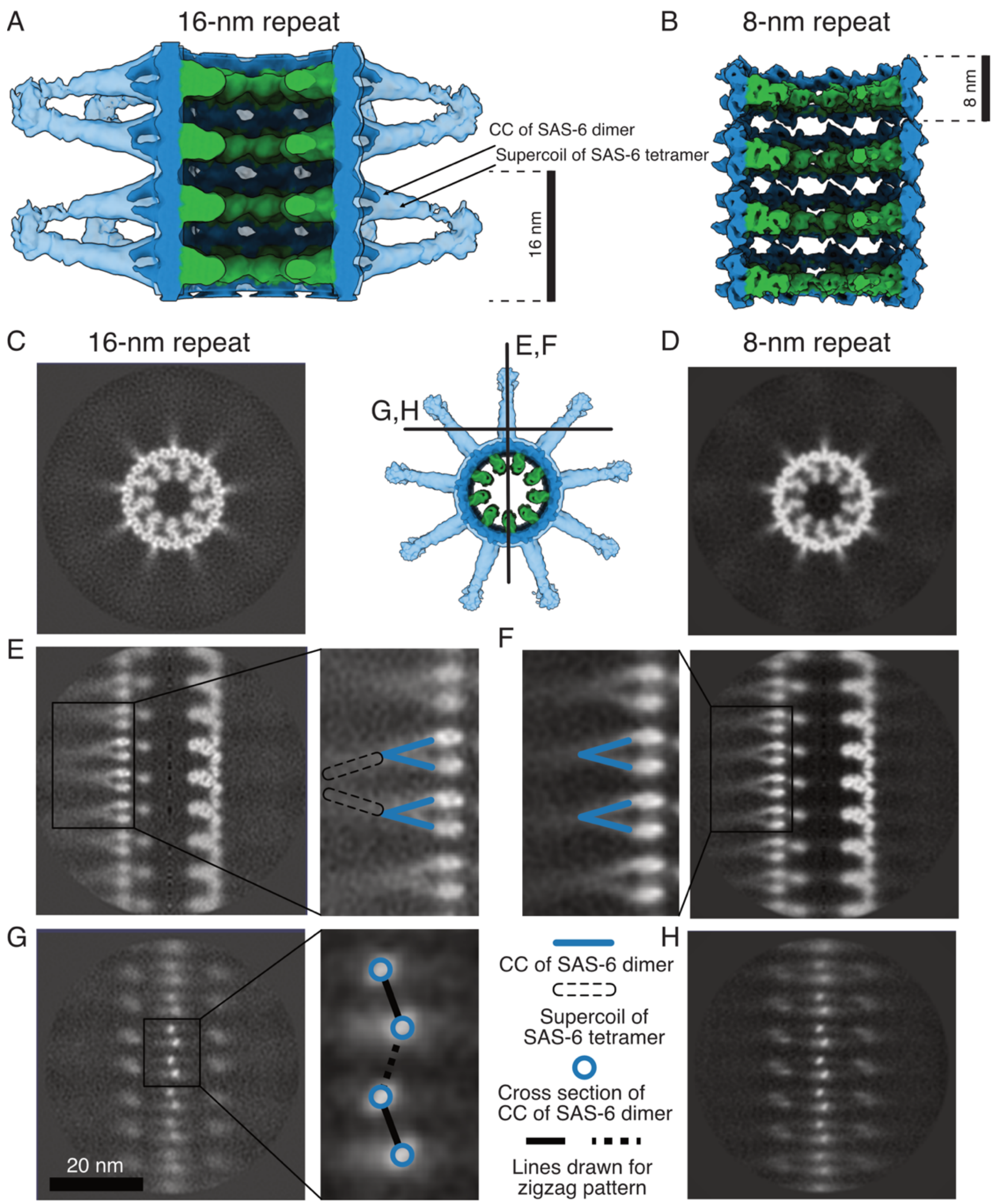


**Fig. S3. Subtomogram averages of the 8-nm and 16-nm CH repeats.** (**A**) Surface rendering of the 16-nm repeat STA map of the CH at 11 Å resolution, displayed within the transparent surface of the 20 Å-filtered CH map (transparent blue) to visualize the prominent spokes. This map is shown at a lower threshold than in Fig. 1D to highlight the segment of the coiled-coil (CC) formed by the SAS-6 dimers that emerges from the density. The transparent blue spoke density is the supercoil of one SAS-6 tetramer, composed of two dimers' coiled-coils. Blue and light blue: SAS-6; green: CID. (**B**) STA map of the 8-nm repeat at 7.6 Å resolution. (**C–H**) Corresponding slices from the 8-nm and 16-nm averaged maps demonstrate consistent structural features across both repeats. (C, D) Cross-sectional views of the CH ring. (E, F) Central slices of the CH-rings. Blue lines indicate their spoke arrangements. (G, H) Slices illustrating the arrangement of SAS-6 dimer CCs and CH rings.

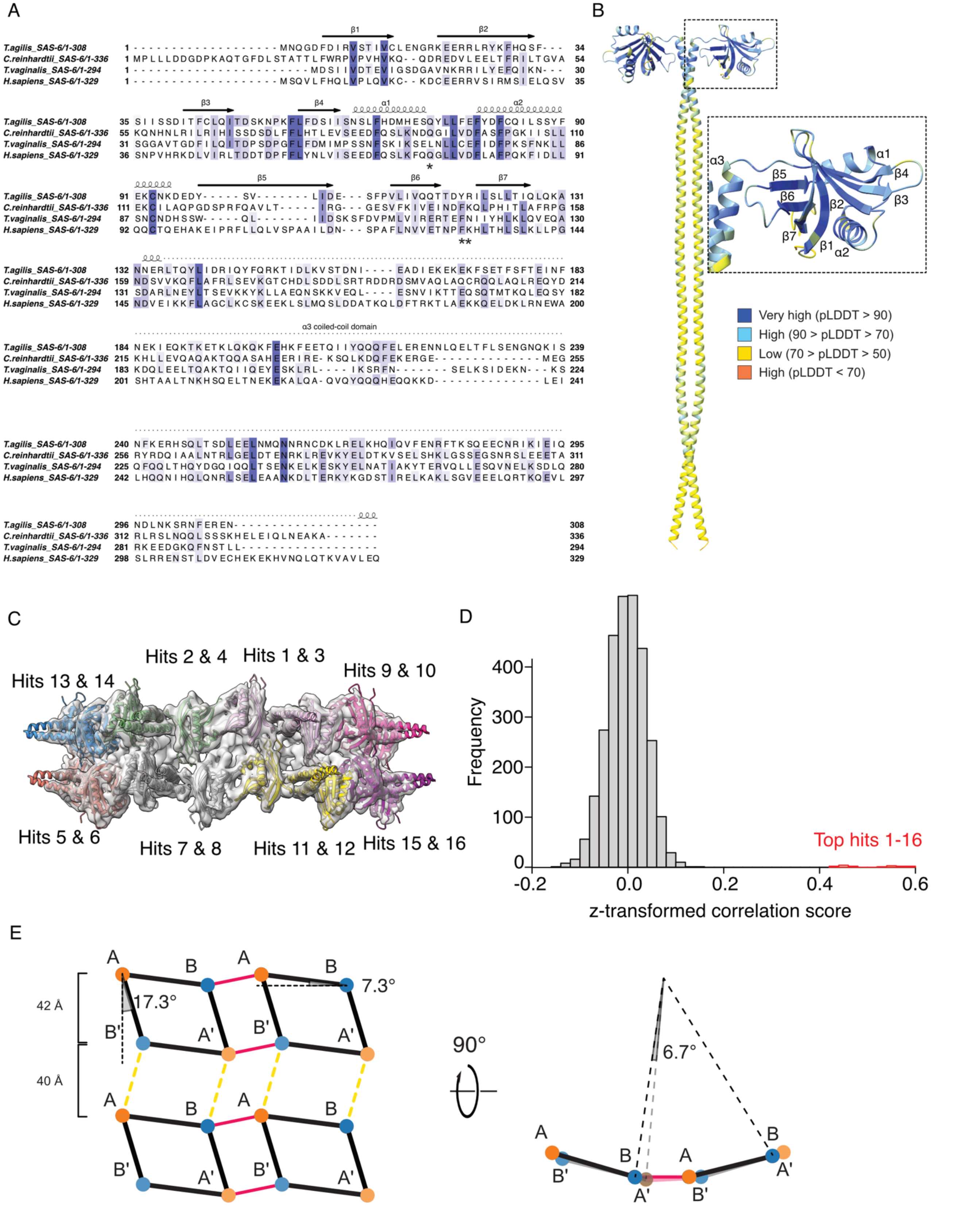


**Fig. S4. Conserved residues in SAS-6 form the CH scaffold. (A)** Multisequence alignment of SAS-$6_{1–308}$ from *Trichonympha agilis* (UniProt: R4WPE9), *Chlamydomonas reinhardtii* (UniProt: A9CQL4), humans (UniProt: Q6UVJ0), and *Trichomonas vaginalis* (UniProt: A2G2L7). More intense color means more conserved. **(B)** AlphaFold3 prediction of the *T. agilis* homodimer colored using predicted local distance difference test (pLDDT) values (interface Predicted TM-score, 0.54; predicted template modelling score, 0.57). **(C)** Top 16 positions of the AlphaFold3 model of the SAS-6 dimer found after fitting with DomainFit (*13*). As the hits and the SAS-6 dimer are C2 symmetric, only one hit is shown. (**D**) The z-transformed correlation scores of the fitting shows that the top hits are well separated from the rest and are true hits. (E) Zigzag stacking ring model of SAS-6 organization with measurement. The individual SAS-6 head domains are drawn in four colors to denote the asymmetric SAS-6 subunits within a tetramer (black parallelogram). Green lines indicate the tetramer-tetramer interaction within the same 8-nm tetramer ring. Dotted yellow lines indicate the stacking interaction between rings. In the right panel, the top view of the zigzag model is viewed from top. The rotational offset between the top AB dimer and the bottom CD dimer is calculated to be 6.7 degrees.

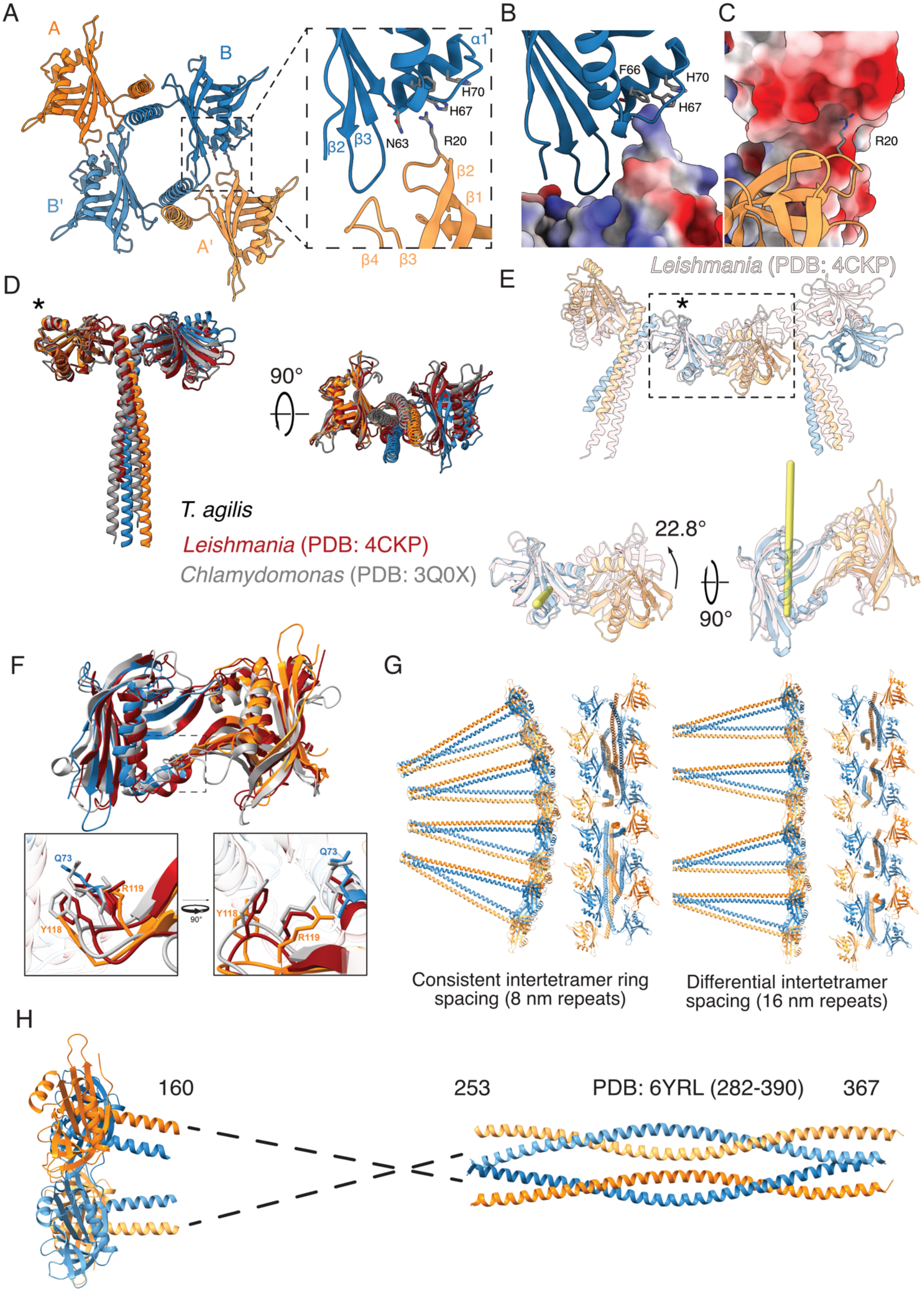
A
A
B
B'
A'
α1
H70
H67
N63
R20
β2
β3
β2
β1
β4
β3
B
F66
H70
H67
C
R20
D
90°
T. agilis
Leishmania (PDB: 4CKP)
Chlamydomonas (PDB: 3Q0X)
E
Leishmania (PDB: 4CKP)
22.8°
90°
F
Q73
R119
Y118
90°
Q73
Y118
R119
G
Consistent intertetramer ring spacing (8 nm repeats)
Differential intertetramer spacing (16 nm repeats)
H
160
253
PDB: 6YRL (282-390)
367

**Fig. S5. Comparison of the *Trichonympha* cartwheel model with existing crystal structures. (A)** Model of the *Trichonympha* tetramerization interface involving helix α1 and loop β2–β3 of chain A′, and loops β1–β2 and β3–β4 of chain B (and the same helix and loops of chains A and B′). **(B)** Electrostatic map of the surface of chain A′ facing chain B. **(C)** Electrostatic map of the surface of chain B facing chain A'. **(D)** Alignment of our TaSAS-6 dimer model (orange and blue, UniProt: R4WPE9) with existing SAS-6 dimer crystal structures from *Chlamydomonas* (gray, PDB: 3Q0X) and *Leishmania* (red, PDB: 4CKP). Symbol (*) denote the N-terminal domain of SAS-6 chain (TaSAS-$1_{1\text{-}130}$) used for alignment. **(E)** Differences in the fits of the *Leishmania* SAS-6 ring and *T. agilis* SAS-6 oligomers reveal the rotation at the ring-forming interface. Symbol (*) denote the N-terminal domain of SAS-6 chain (TaSAS-$6_{1\text{-}130}$) used for superimposition. The SAS-6 N-terminal domain of the neighboring SAS-6 dimer rotates 22.8° around an axis (yellow line) almost parallel to the CH axis. **(F)** Conservation of key inter-tetramer interface residues and the hydrophobic pocket across SAS-6 homologs. **(G)** Consistent inter-tetramer spacing creates a vertical curvature along the cartwheel length, while gaps between two SAS-6 tetramer layers enable straight assembly. **(H)** Alignment of tetramerized *T. agilis* SAS-6 N-terminal domains tetramer with the coiled coil domains of the *Chlamydomonas* SAS-6 homolog Bld12 (PDB: 6YRL; residues 282–390, equivalent to TaSAS-$6_{253\text{–}367}$.

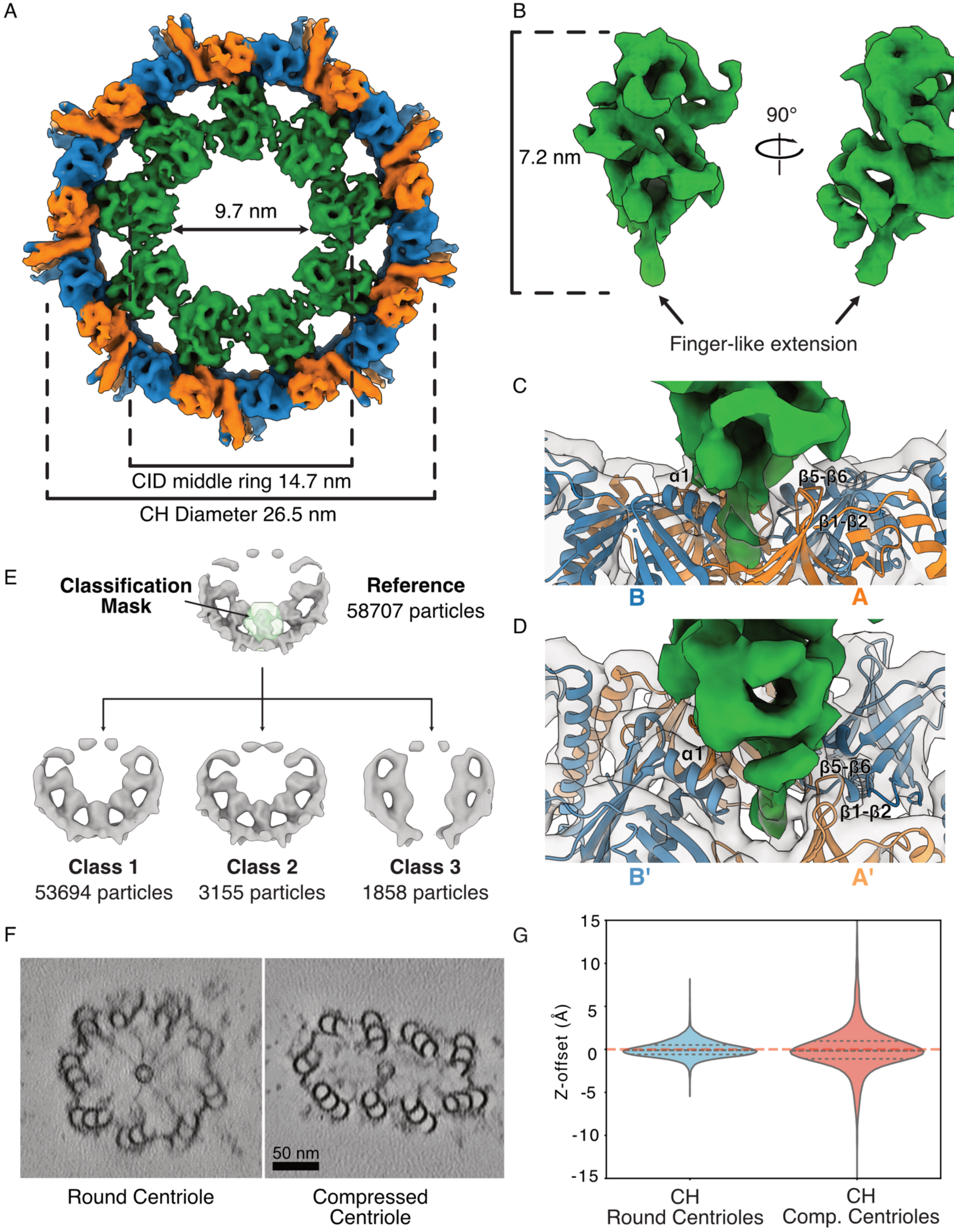


**Fig. S6: The CID forms an inner ring inside the CH. (A)** Color rendering of one CH ring at a lower threshold than in Fig. 3A, showing potential linkages between the CID densities. Viewed from the distal side. Measurement of indicating the CH ring diameter, CID middle ring diameter, and conduit diameter. **(B)** A segmented CID density in two orientations. **(C)** Interactions between the CID and chains A and B. **(D)** Interactions between the CID and chains A′ and B′. (**E**) Classification of CID densities from the subunits of 8-nm ring into 3 classes. (**F**) Views of 14 nm-thick tomographic section of round and compressed centrioles showing that the CH keep its round shape. **(G)** Combined violin plots show the distribution of vertical deviations, with lines marking the median and interquartile ranges. The mean absolute Z-offset (|Z|) rose from 0.69 ± 0.60 Å to 1.67 ± 1.93 Å, indicating a significant loss of planarity under compression (p value = $3.3667 \times 10^{-255}$).

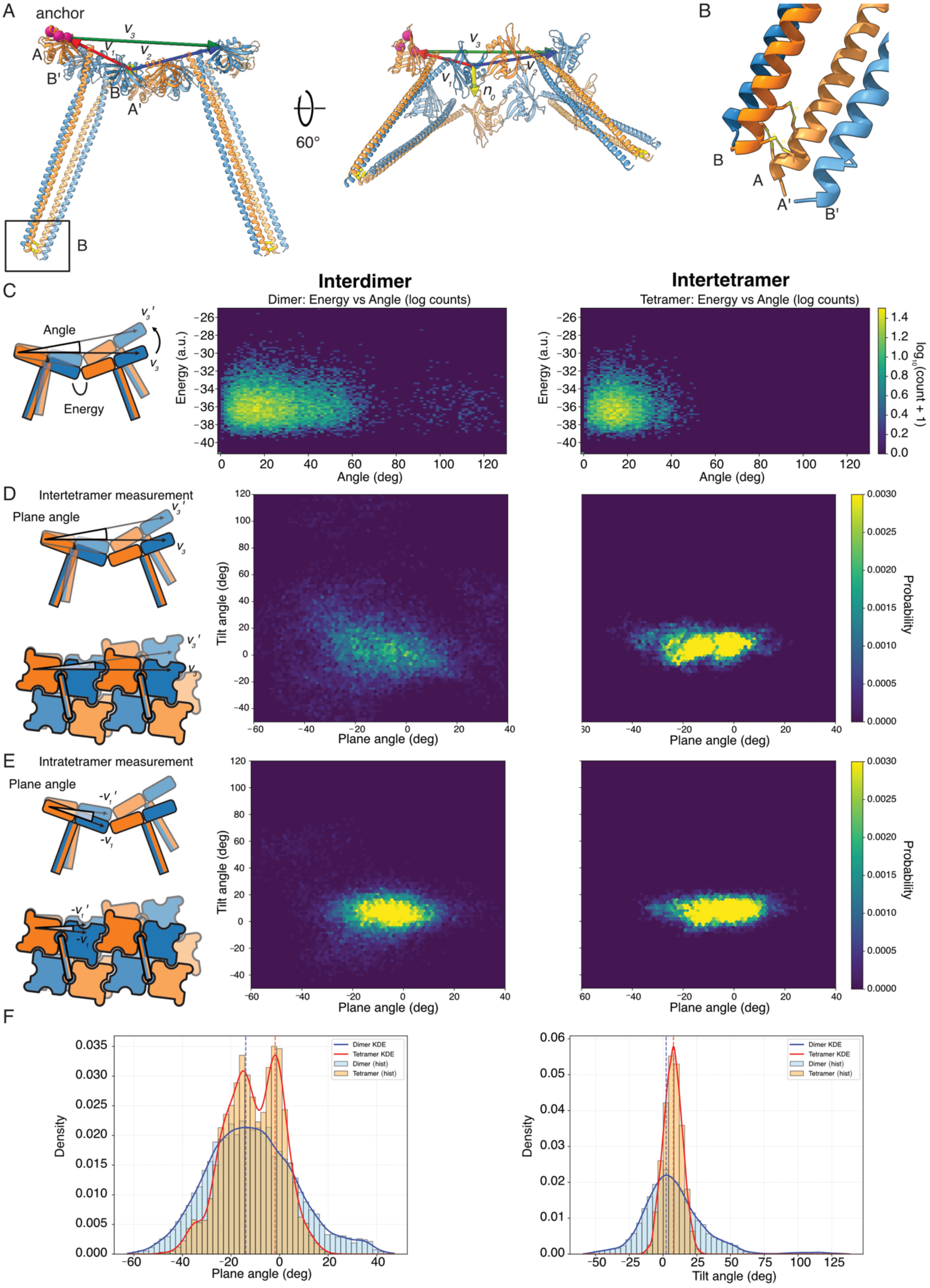
A
anchor
B
C
Interdimer
Dimer: Energy vs Angle (log counts)
Intertetramer
Tetramer: Energy vs Angle (log counts)
Angle
Energy
Energy (a.u.)
Angle (deg)
log10(count + 1)
D
Intertetramer measurement
Plane angle
Tilt angle (deg)
Plane angle (deg)
Probability
E
Intratetramer measurement
F
Density
Dimer KDE
Tetramer KDE
Dimer (hist)
Tetramer (hist)
Tilt angle (deg)
60°

**Fig. S7**. **Coarse-grained molecular dynamics of SAS-6 interactions (A)** The inter-tetramer model ($SAS\text{-}6_{1\text{-}250}$) used in the simulation. Vectors $v_1$, $v_2$, $v_3$ define the geometry for plane and tilt angle calculations; $n_0$ is the normal vector to the plane formed by $v_1$ and $v_2$ and was used to compute the tilt angle. Anchor points (the Cα atoms of N63, Q73, and Q127) are shown as purple spheres. **(B)** Cysteine substitutions at residues 235, 238, and 239 of chains B and B′ to promote disulfide bond formation between the coiled coil regions, mimicking the structural effect of supercoil formation. **(C)** Interaction energies of the dimer–dimer and tetramer–tetramer assemblies at the inter-unit interface. **(D)** Tilt and plane angle distributions of the interdimer and inter-tetramer interactions, calculated based on deviations relative to $v_3$. **(E)** A control analysis measuring tilt and plane angle variations between the two SAS-6 molecules within the same dimer, demonstrating that the flexibility observed in (D) arises from interdimer or inter-tetramer interactions, as the intradimer geometry is relatively rigid. **(F)** Histogram and kernel density estimation plots of the plane and tilt angles from the control analysis in (E).
